# Changing microbial activities during low salinity acclimation in the brown alga *Ectocarpus subulatus*

**DOI:** 10.1101/2021.04.13.439635

**Authors:** Hetty KleinJan, Gianmaria Caliafano, Méziane Aite, Enora Fremy, Clémence Frioux, Elham Karimi, Erwan Corre, Thomas Wichard, Anne Siegel, Catherine Boyen, Simon M. Dittami

## Abstract

*Ectocarpus subulatus* is one of the few brown algae found in river habitats. Its ability to tolerate freshwater is due, in part, to its uncultivated microbiome. We investigated this phenomenon by modifying the microbiome of laboratory-grown *E. subulatus* using mild antibiotic treatments, which affected its ability to grow in low salinity. The acclimation to low salinity of fresh water-tolerant and intolerant holobionts was then compared. Salinity had a significant impact on bacterial gene expression as well as the expression of algae- and bacteria-associated viruses in all holobionts, albeit in different ways for each holobiont. On the other hand, gene expression of the algal host and metabolite profiles were affected almost exclusively in the fresh water intolerant holobiont. We found no evidence of bacterial protein production that would directly improve algal stress tolerance. However, we identified vitamin K synthesis as one possible bacterial service missing specifically in the fresh water-intolerant holobiont in low salinity.

We also noticed an increase in bacterial transcriptomic activity and the induction of microbial genes involved in the biosynthesis of the autoinducer AI-1, a compound that regulates quorum sensing. This could have caused a shift in bacterial behavior in the intolerant holobiont, resulting in virulence or dysbiosis.

**Originality-Significance Statement:** The importance of symbiotic microbes for the health and stress resistance of multicellular eukaryotes is widely acknowledged, but understanding the mechanisms underlying these interactions is challenging. They are especially difficult to separate in systems with one or more uncultivable components. We bridge the gap between fully controlled, cultivable model systems and purely environmental studies through the use of a multi-omics approach and metabolic models on experimentally modified “holobiont” systems. This allows us to generate two promising working hypotheses on the mechanisms by which uncultivated bacteria influence their brown algal host’s fresh water tolerance.

## Introduction

Brown algae are multicellular members of the stramenopile lineage and frequently form the dominant vegetation in intertidal zones of temperate marine coastal ecosystems (Wahl *et al.*, 2015). They are important both as ecosystem engineers and increasingly being exploited as a food source (Food and Agriculture Organization of the United Nations, 2016), and for the production of alginate (McHugh, 2003) or other high-value compounds (Milledge *et al.*, 2016; Silva *et al.*, 2020). Like virtually all macroorganisms, brown algae have formed tight relationships with their associated microbiota, which may provide them with vitamins, phytohormones, protection against disease or fouling, etc. (Goecke *et al.*, 2010; Wahl *et al.*, 2012; Egan and Gardiner, 2016).

*Ectocarpus* is a cosmopolitan genus of small filamentous brown algae that is easy to cultivate in the laboratory but closely related to large kelp-forming brown algal species, at the evolutionary level. The *Ectocarpus* sp. strain Ec32 has been established as one of the key model systems to study brown algal biology (Charrier *et al.*, 2008), and its genome was the first brown algal genome to be published (Cock *et al.*, 2010). *Ectocarpus* is furthermore a model to study algal bacterial interactions, with studies demonstrating, for instance, its reliance on bacteria-produced cytokinins, which serve as morphogens for the alga (Pedersén, 1968, 1973; Tapia *et al.*, 2016). More recently, metabolic complementarity has been successfully used in this system, highlighting potentially beneficial metabolic host-microbe interactions (Dittami *et al.*, 2014; Burgunter-Delamare *et al.*, 2020).

One species of *Ectocarpus*, *Ectocarpus subulatus*, is of particular interest, as it has been described in river habitats (West and Kraft, 1996; Dittami, Peters, *et al.*, 2020), yet its capacity to tolerate freshwater medium in the laboratory seems to be conditioned by the presence of a specific microbiota (Dittami *et al.*, 2016). We speculate that this dependence on bacteria may be related to (i) the direct production of compounds such as osmolytes or chaperones that enhance algal low salinity tolerance, (ii) general bacterial services that are required for algal growth regardless of the salinity, (iii) protective functions of some bacteria to prevent phenomena of dysbiosis, or (iv) a combination of these effects. However, despite extensive cultivation efforts (KleinJan *et al.*, 2017) followed by co-culture experiments (personal data), neither the bacteria responsible for this phenomenon nor the underlying bacterial functions have been identified so far, possibly because (part of) the bacteria responsible are found within the uncultivable part of the microbiome.

The role of *Ectocarpus*-associated bacteria in the fresh water response is investigated in this study using a novel approach. Rather than starting from a collection of cultured bacteria, we worked with algal holobionts that had been treated with various antibiotic combinations. These antibiotics were powerful enough to change the algal microbiome, but not completely eliminate it. Several of the treated algae were able to survive in freshwater. We then used a combination of metagenomics, meta-transcriptomics, and metabolomics to investigate three algal holobionts, each with its own microbiome and response to fresh water. It allowed us to develop testable hypotheses about how these holobionts respond to low salinity and, in particular, how changes in bacterial composition and activity correlate with the alga’s acclimation capacity.

## Material and methods

### Preparation of algal holobionts with modified microbiomes

Starter cultures of *Ectocarpus subulatus* freshwater strain (EC371, accession CCAP 1310/196, West & Kraft, 1996) were grown in 90 mm Petri dishes in natural seawater (NSW; salinity 35, collected in Roscoff 48°46’40″N, 3°56’15″W, 0.45 μm filtered, autoclaved at 120°C for 20 min), enriched with Provasoli nutrients (2 mg·L^−1^ Na_2_EDTA, 2.24 mg·L^−1^ H_3_BO_3_, 240 μg·L^−1^ MnSO_4_·H_2_O, 44 μg·L^−1^ ZnSO_4_·7 H_2_O, 10 μg·L^−1^ CoSO_4_·7H_2_O, 0.7 mg·L^−1^ FeEDTA, 0.6 mg·L^−1^ Na_2_EDTA, 4 mg·L^−1^ Na_2_ ß-glycerol PO_4_·5H_2_O, 35 mg·L^−1^ NaNO_3_, 0.875 μg·L^−1^ vitamin B12, 40 μg·L^−1^ thiamine, 4 μg·L^−1^ biotin; Starr and Zeikus 1993). They were kept at 13°C with a 12h dark-light cycle (photon flux density 20 μmol m^−2^·s^−1^). Starter cultures were then treated with different antibiotics, and their capacity to grow in diluted natural seawater medium (5% NSW, 95% distilled water) was assessed (Supplementary Table S1). Three “holobionts” were selected for the final experiments (Table 1). Two of them (H1, H2) were based on short (3 days) treatments with antibiotics solutions and were capable of growing in diluted natural seawater medium. The last one (H3) was subjected to 5 weeks of antibiotic treatment on Petri dishes and was no longer fresh water-tolerant. After recovery of two weeks (H1, H2) to five months (H3) in antibiotic-free 100% NSW, cultures were transferred to 10 L culture flasks, grown for approximately two months to obtain sufficient biomass, and then split into ten replicate cultures in 2 L culture flasks (each replicate with identical microbial communities). To ensure that the holobionts differed in their microbial community composition, and before acclimation experiments, two replicates samples of each culture (technical replicates) were used for 16S rRNA gene metabarcoding. The experimental setup is shown in Figure 1.

**Table 1:**
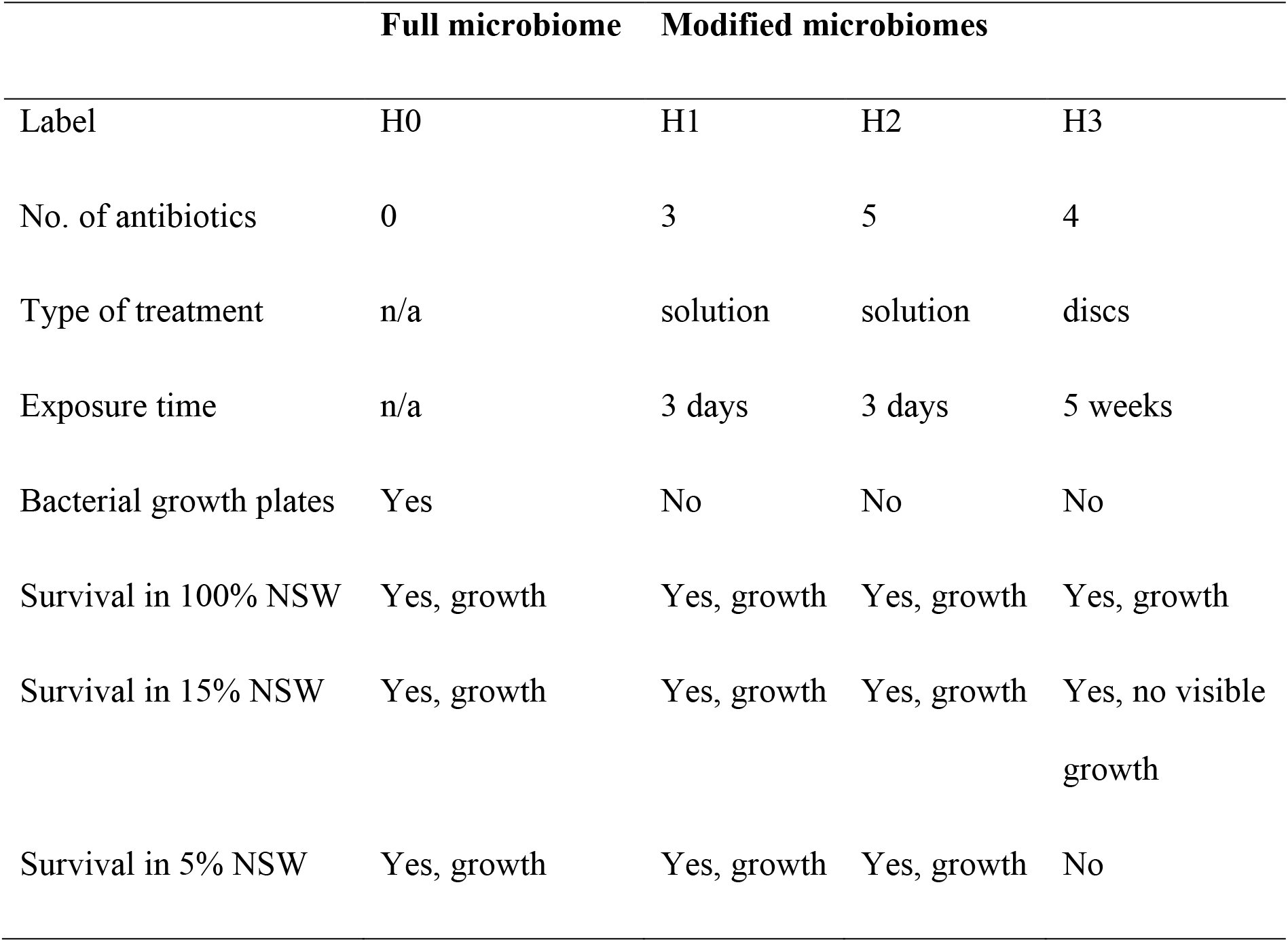
Different algal holobionts used for the metatranscriptomics/metabolomics experiment; Holobiont H1: treated with rifampicin, penicillin and neomycin (each 100 μg/ml); Holobiont H2: idem, plus streptomycin (25 μg/ml) and chloramphenicol (5 μg/ml); Holobiont H3: treated with penicillin (12000UI), Chloramphenicol (0.75 μg/ml), Polymyxin B (0.75 μg/ml), Neomycin (0.9 μg/ml). The untreated holobiont (H0) was only used for 16S rRNA gene metabarcoding, not for metatranscriptomics/metabolomics. Algae were considered “alive” as long as their pigmentation was visible, even in the absence of visible growth.

**Figure 1:**
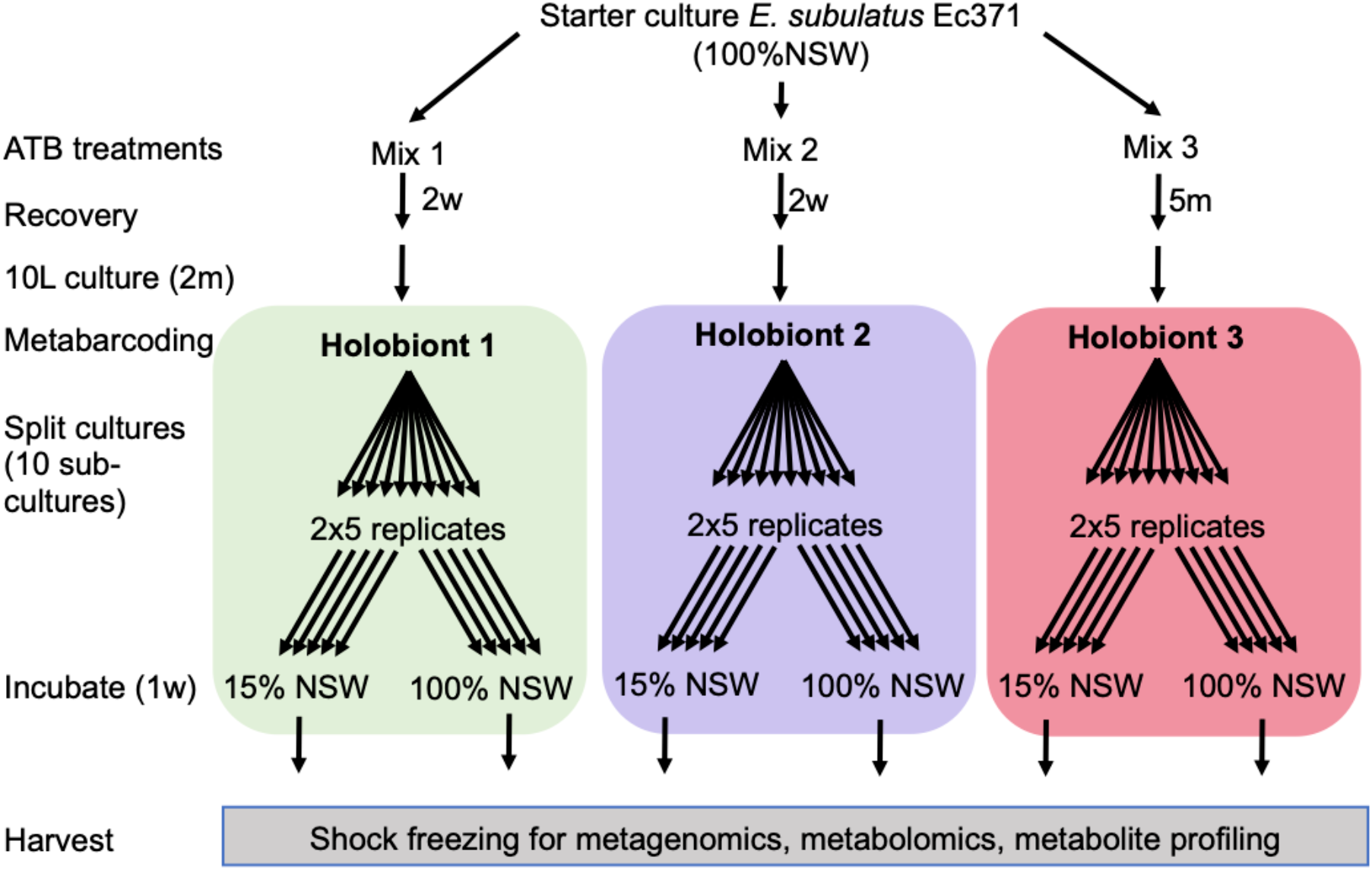
Overview of the experimental setup. All conditions were derived from the same starter culture, but antibiotic (ATB) treatments were carried out with three different ATB mixes (see methods) leaving hosts with different microbial communities, termed “holobionts 1-3”. The low-salinity response of each of these holobionts was then examined with 5 replicates. 100% NSW=natural seawater, 15% NSW=15% NSW in distilled H2O (volume/volume), m=month(s), w=week(s).

### 16S rRNA gene metabarcoding

16S rRNA metabarcoding was carried out as previously described (KleinJan *et al.*, 2017). Briefly, total DNA was isolated (NucleoSpin Plant II, Machery-Nagel; standard protocol) from snap-frozen tissue and purified with Clontech CHROMA SPINTM-1000+DEPC-H2O columns. The V3–V4 region of the 16S rRNA gene was amplified and sequenced with Illumina MiSeq technology by MWG Eurofins Biotech (Ebersberg, Germany) using their proprietary protocol and yielding 1,859,076 reads. After quality trimming using the FASTX Toolkit (quality threshold 25; minimum read length 200), data were analyzed with Mothur (V.1.38.0) according to the MiSeq Standard Operating Procedures (Kozich *et al.*, 2013). Sequences were aligned to the non-redundant Silva SSU reference database version 123 (Quast *et al.*, 2013), chimeric sequences removed using Uchime (Edgar *et al.*, 2011), clustered into operational taxonomic units (OTUs) at a 97% identity level, and classified taxonomically (Wang *et al.*, 2007).

### Acclimation experiments

Acclimation experiments were carried out to elucidate differences in the response of the three holobionts to low salinity. Because natural seawater medium diluted to 5% NSW was lethal to the non-tolerant holobionts (H3) and based on preliminary screening experiments, we opted for a final concentration of 15% NSW and 85% distilled water enriched with Provasoli nutrients. At 15% NSW, growth in the non-tolerant holobiont (H3) was inhibited, but the condition was not lethal. Five replicate cultures of each holobiont (prepared as described above) were transferred from 100% NSW to 15% NSW, and the other five replicates were transferred to fresh 100% NSW as a control. After one-week, algal tissue was harvested using UV-sterilized coffee filters. Excess water was removed by drying with a clean paper towel, tissues snap-frozen in liquid nitrogen, and stored at −80 °C until further processing.

### Metagenome and metatranscriptome sequencing

Approximately 50 mg (fresh weight) of algal tissue were ground in liquid nitrogen and sterilized sand using a pestle and mortar. Nucleic acids were extracted as previously described by Le Bail *et al.* (2008) using a CTAB-based lysis buffer and phenol-chloroform purification. RNA and DNA were separated after precipitation with LiCl (0.25V, 12M, overnight, −20°C); the RNA was resuspended in 300 μl RNase free and purified with 1 volume of phenol: chloroform (1:1, pH 4.3, 20 min., 4°C) and twice with one volume of chloroform. It was then precipitated (0.1V 3M sodium acetate, 3V 100% ethanol, 2h at −80°C), washed with ice-cold 70% ethanol, and finally resuspended in 15 μl RNase free water. Ribosomal rRNA molecules were depleted from the RNA extracts using the RiboMinus™ Plant Kit for RNA-Seq (ThermoFisher Scientific, Waltham, MA, USA). This allowed to selectively remove abundant nuclear, mitochondrial, and chloroplast rRNAs of the algal host. Besides, 1 μl of the bacterial probe (RiboMinus™ Transcriptome Isolation Kit for Yeast and Bacteria, ThermoFisher Scientific) was added in the last five minutes of hybridization to remove bacterial ribosomal RNA. The RNA was concentrated (RiboMinusTM Concentration module, ThermoFisher Scientific) before checking the quality with a bioanalyzer (Agilent, Santa Clara, CA, USA). Library preparation (TruSeq Stranded mRNA Library Prep kit, Illumina) and sequencing (five lanes, 150bp read length, paired-end, Illumina HiSeq 3000, GeT PlaGe Genotoul, Auzeville, France) were carried out for four of the five replicates per condition. The fifth replicate was kept as a backup.

DNA was extracted from the supernatant of LiCl precipitation. It was precipitated with one volume of isopropanol, resuspended in 300 μL DNA-free water, purified once with one volume of phenol:chloroform:isoamylic alcohol (25:24:1; pH 8), and twice with one volume of chloroform. Finally, the DNA was precipitated (3 volumes of 100% EtOH + 0.1 volumes of 3M sodium acetate) and resuspended in 50 μL of molecular biology grade water. DNA extracts from all samples were pooled (same concentration of each sample), and the pooled DNA was purified using cesium chloride gradient centrifugation (Le Bail *et al.*, 2008). A library was constructed using the Illumina TruSeq DNA Nano kit and was sequenced on four lanes of Illumina HiSeq 3000 (150bp read length, paired-end, GeT PlaGe Genotoul platform). DNA and RNA sequencing data were deposited at the European Nucleotide Archive (ENA) under project accession PRJEB43393.

### Metagenome analyses

Raw sequencing reads were quality-trimmed with Trimmomatic (version 0.36, minimal Phred score: 20, minimal read length: 36 nucleotides) and aligned to the *E. subulatus* Bft15b reference genome (Dittami, Corre, *et al.*, 2020) using STAR aligner version 2.6.0a (Dobin *et al.*, 2013) to remove algal reads. Non-aligning (*i.e.* bacterial) reads were then assembled using MetaSPAdes (Nurk *et al.*, 2017). The resulting contigs were filtered by length (>500 bp), and the remaining *Ectocarpus* contigs were removed with Taxoblast version 1.21 (Dittami and Corre, 2017). Metagenomic binning was carried out using Anvi’o version 4.0 according to the “Anvi’o User Tutorial for Metagenomic Workflow” (Eren *et al.*, 2015): Raw metagenome reads were mapped against the contigs database using BWA-MEM (version 0.7.15), and taxonomy was assigned to the contigs using Centrifuge (Kim *et al.*, 2016) and the NCBI nucleotide non-redundant database (nt_2018_3_3). Finally, contigs were clustered according to GC content and coverage, and the Anvi’o interactive interface was used for manual curation of the bins. The quality of the metagenomes was assessed based on the abundance of single-copy core genes (SCGs; Campbell et al., 2013). One bin containing essentially *Ectocarpus* reads that had been missed during the previous cleaning steps was manually removed at this stage. The remaining bins were bacterial and annotated with Prokka v1.13 (Seemann, 2014).

Metabolic networks were reconstructed using Pathway Tools version 20.5 (Karp *et al.*, 2016) and the scripts included in the AuReMe pipeline (Aite *et al.*, 2018). This dataset served as a backbone for the analysis of bacterial gene expression from the metatranscriptomic data.

### (Meta)transcriptome analyses

Ribosomal RNA reads that remained despite the RiboMinus treatment were removed *in silico* using SortMeRNA version 2.1. After quality trimming (Trimmomatic 0.36, minimal Phred score: 20, minimal read length: 36 nucleotides), reads were mapped first to the *E. subulatus* Bft15b reference genome for algal gene expression analysis and the remaining reads to the metagenomic bins generated as described above using STAR aligner version 2.6.0a. The ratio of bacterial to algal mRNA was compared across samples using a one-way ANOVA followed by Tukey’s HSD test carried out using Past v4 (Hammer *et al.*, 2001). Then, both the algal transcriptome and the bacterial metatranscriptome were analyzed separately. In both cases, DESeq2 (Love *et al.*, 2014) was used for principal component analysis (PCA, rlog-transformed data) and for the detection of significantly differentially expressed genes (DEGs). However, for the bacterial metatranscriptome, read coverage was insufficient to carry out differential gene expression analysis on a bin per bin basis. Therefore, we summed up the bacterial expression data associated with the same metabolic reaction across the different metabolic networks. This overall transcription of a metabolic function within the entire microbiome was used to identify differentially expressed microbial reactions in the same way as the algal DEGs.

For both the algal transcriptome and the bacterial metabolic reactions, the following comparisons were carried out. First, each of the three holobionts (H1, H2, H3) was analyzed individually to determine genes and reactions that were differentially expressed in 15% NSW compared to 100% NSW. Then, the same conditions of the two freshwater-tolerant holobionts were grouped (i.e. H1-100% and H2-100% were grouped as well as H1-15% and H2-15%), and these newly formed groups were compared to the holobiont that was not able to grow in freshwater (H3-100% and H3-15%). For this comparison, the statistical design considered both factors (holobiont and salinity) and the interaction term. All genes or reactions with an adjusted p-value <0.05 and a fold change in expression > 1.5 were considered significantly differentially expressed. Gene set enrichment analyses were performed for sets of differentially expressed genes using the Fisher’s exact module within Blast2Go (Version 4.1.9; 2-tailed test; FDR < 0.05; Götz *et al.* 2008). Metabolic reactions were associated with metabolic pathways according to the MetaCyc database v20.5, and pathways with >50% of differentially expressed metabolic reactions were further examined.

For the bacterial metatranscriptomes, in addition to differentially expressed metabolic reactions, we also determined the overall “transcriptomic activity” of each bin in each condition. To this means, read pair counts were summarized for all genes of the same bin and normalized by the total number of mapping read pairs per sample. The resulting matrix was used as input for hierarchical clustering (distance: correlation; method: average) with ClustVis (Metsalu and Vilo, 2015).

Finally, viral reads present in the metatranscriptomes were analyzed using the Metagenomic RAST (MG-RAST) pipeline (Meyer *et al.*, 2008). Sequencing reads were annotated using the best-hit annotation tool against the RefSeq database (Pruitt *et al.*, 2007). Taxonomic affiliations were generated at the family level if possible, and the BLAST parameters were set to the default settings of MG-RAST (15aa minimal alignment and 1E−05 e-value threshold). The extracted number of reads for viruses was Hellinger-normalized and used as input for ANOVA and PCA analyses using STAMP v2.1.3 (Parks *et al.*, 2014).

### Metabolite profiling by gas-chromatography

For metabolite profiling, 10 mg of freeze-dried ground tissue was lysed (TissueLyser, 2*30s, 30 Mhz, Qiagen, Hilden, Germany) and subsequently used for extraction of metabolites with 1 ml of 100% methanol (Dittami *et al.*, 2012). After vortexing, 5 μl of ribitol (4 mM in H_2_O, >99%, Sigma-Aldrich, Munich, Germany) were added to each sample as an internal standard before sonication (10 min, at room temperature; RT). After 15 minutes of centrifugation (30,000 g, 4 °C) the supernatant was recovered, evaporated under vacuum overnight, derivatized in 50 μl methoxymation solution (20 mg/ml in pyridine), and incubated at 60 °C for 1h and afterward at RT overnight. The samples were silylated for 1h at 40°C in 50 μl MSTFA (1 ml + 40 μl retention index mix) and centrifuged (6 min, 2500 rpm) to pellet the precipitate (Alsufyani *et al.*, 2017). The supernatant was analyzed with a 6890N gas chromatograph, equipped with a 7683B autosampler (Agilent), a glass liner (Agilent, 4 × 6.3 × 78.5 mm), and a DB-5MS column (Agilent, 30 m × 0.25 mm × 0.25 μm), coupled to a Micromass GCT Premier™ (Waters®) mass spectrometer. The gas chromatograph was operated with helium as a mobile phase, split 10, and 250°C injector temperature. The initial oven temperature was 60°C ramping to 310°C at a rate of 15°C per min. The mass spectrometer was used with a source temperature at 300°C and dynamic range extension mode. The resolution was > 6,000 FWHM at *m/z* 501.97. After randomization, samples were measured twice to obtain two technical replicates. Solvent blanks were prepared and measured in parallel.

### Analysis of metabolite data

Raw files were directly converted to the netCDF format using the DataBridge tool within the MassLynx software (Waters, version 4.1), and the chromatograms were then processed with the function metaMS.runGC (version 1.0) provided within Workflow4Metabolomics (W4M) (Giacomoni *et al.*, 2015). The metaMS package (Wehrens *et al.*, 2014) was used to identify chromatographic peaks with the standard functions provided by XCMS. Then, the CAMERA package was used to cluster masses with similar retention times (Kuhl *et al.*, 2012). These co-eluting masses or ‘pseudospectra’ were summarized into a final feature table in the MSP format, a format that can be used to search in spectral databases. A detailed list of settings can be found in Supplementary Table S2. The resulting matrix of 689 features (pseudospectra) was manually processed. To remove any contaminant signals from the matrix, each peak’s maximum value among all blanks was multiplied by three and subtracted from the remaining samples. Variables with less than two samples with intensities above zero and redundant ions (isotopes) were removed. The filtered datasets were then re-imported into W4M for statistical analysis. Quality assessment of the data confirmed that there were no outliers and there was no signal drift. Data were normalized by dry weight followed by log10-scaling and a student *t*-test was used to detect metabolites that were significantly different (adjusted p-value < 0.05) in each holobiont during the shift from 100% NSW to 15% NSW, and between holobionts H1/H2 and holobiont H3 in the 100% NSW condition. Finally, the spectra of each significant feature were compared to the GOLM libraries (Hummel *et al.*, 2010) and an in-house library (Kuhlisch *et al.*, 2018) for annotation using NIST MS Search (version 2.0). Features with a reverse match score (R) ≥800 were annotated, for 700≤R<800 features were annotated but labeled with an additional “?”, and for 600≤R<700 they were labeled with “??”. Clustering was carried out using ClustVis as described above for the metatranscriptome data.

## Results

### Sequencing data

Illumina sequencing resulted in a total number of 2.9 billion metagenomic reads and 3.2 billion RNAseq reads (average of 2.7 million per sample). Roughly half of the metagenome reads mapped with the algal genome, and the other half was considered bacterial. For the RNAseq data, despite *in vitro* ribodepletion, on average, 81% of reads corresponded to ribosomal sequences, and the remaining reads mapped to the alga (both nuclear 2-11% and organellar reads 1-21%). Only 0.4 to 12% (3% on average) of the total reads did not map to the algae (see Supplementary Table S3 for details) and were considered bacterial or viral. The ratio of bacterial to algal mRNA varied significantly according to treatment (one-way ANOVA, p < 0.001) and was highest in H3 in low salinity, which differed significantly from all other treatments according to a Tukey HSD test (p < 0.001, Figure 2).

**Figure 2:**
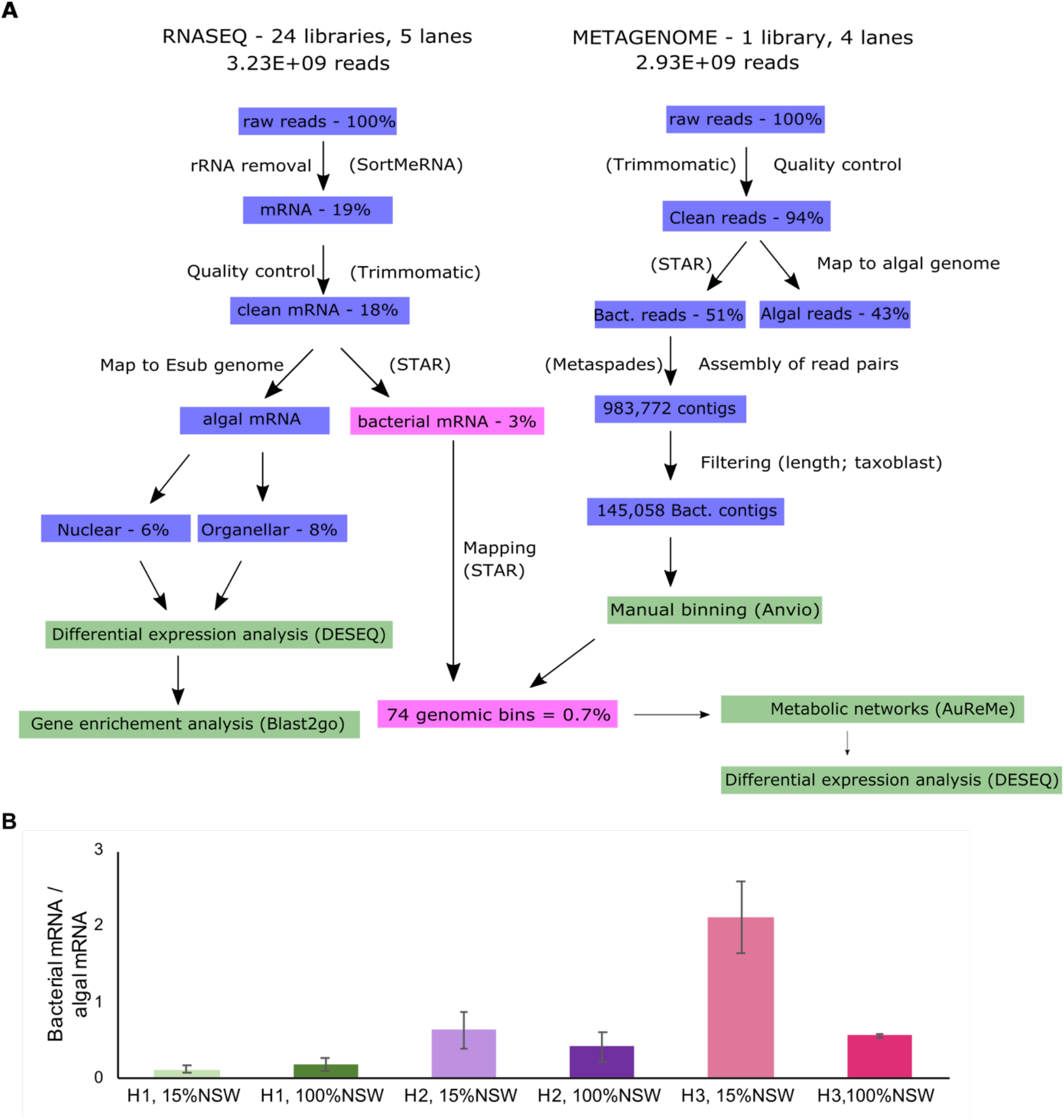
A) Overview of metatranscriptomic and metagenomic data obtained and the analysis pipeline. The percentages correspond to the percentage of total raw reads at the start. B) ratio of bacterial to algal mRNA in the different holobionts/conditions (mean of 4 replicates ± SD). H3-15% is significantly different from all other conditions (p<0.001 Tukey HSD test.)

### The bacterial metagenome

Metagenome sequencing of pooled DNA of all samples was carried out to generate a reference for the analysis of bacterial gene expression. The assembly of metagenome reads not mapping to the algal genome resulted in a total of 145,058 contigs corresponding to 332 Mbp of sequence information. Anvi’o binning of these contigs yielded 73 bins (Figure 3), as well as one bin that was created artificially to regroup all contigs that did not fall into any other well-defined bin (19 Mbp of sequence data). Thirty-five metagenomic bins had a completeness ≥ 90% (categorized as “full”). Four of those were ≥10% redundant (Bin 29: 18%, Bin 61: 18%, Bin 74: 16%, Bin 42: 21%). Thirty-eight bins were ≤ 90% complete and categorized as “partial”. Most bins were taxonomically assigned to *Proteobacteria* (53), followed by *Bacteroidetes* (11) *Planctomycetes* (3), *Actinobacteria* (3), and unclassified bacteria (3). Among the *Proteobacteria*, *Alphaproteobacteria*, specifically *Rhizobiales* (12) and *Rhodobacterales* (15), were the most abundant. *Bacteroidetes* were comprised of six *Flavobacteria* and one *Cytophaga*. A complete overview of the bacterial bins and assembly statistics can be found in Supplementary Table S4. Each of these genomic bins was annotated, and metabolic networks were created. The merged metabolic network of all bacterial bins comprised 3,957 different metabolic reactions (Supplementary Table S5).

**Figure 3:**
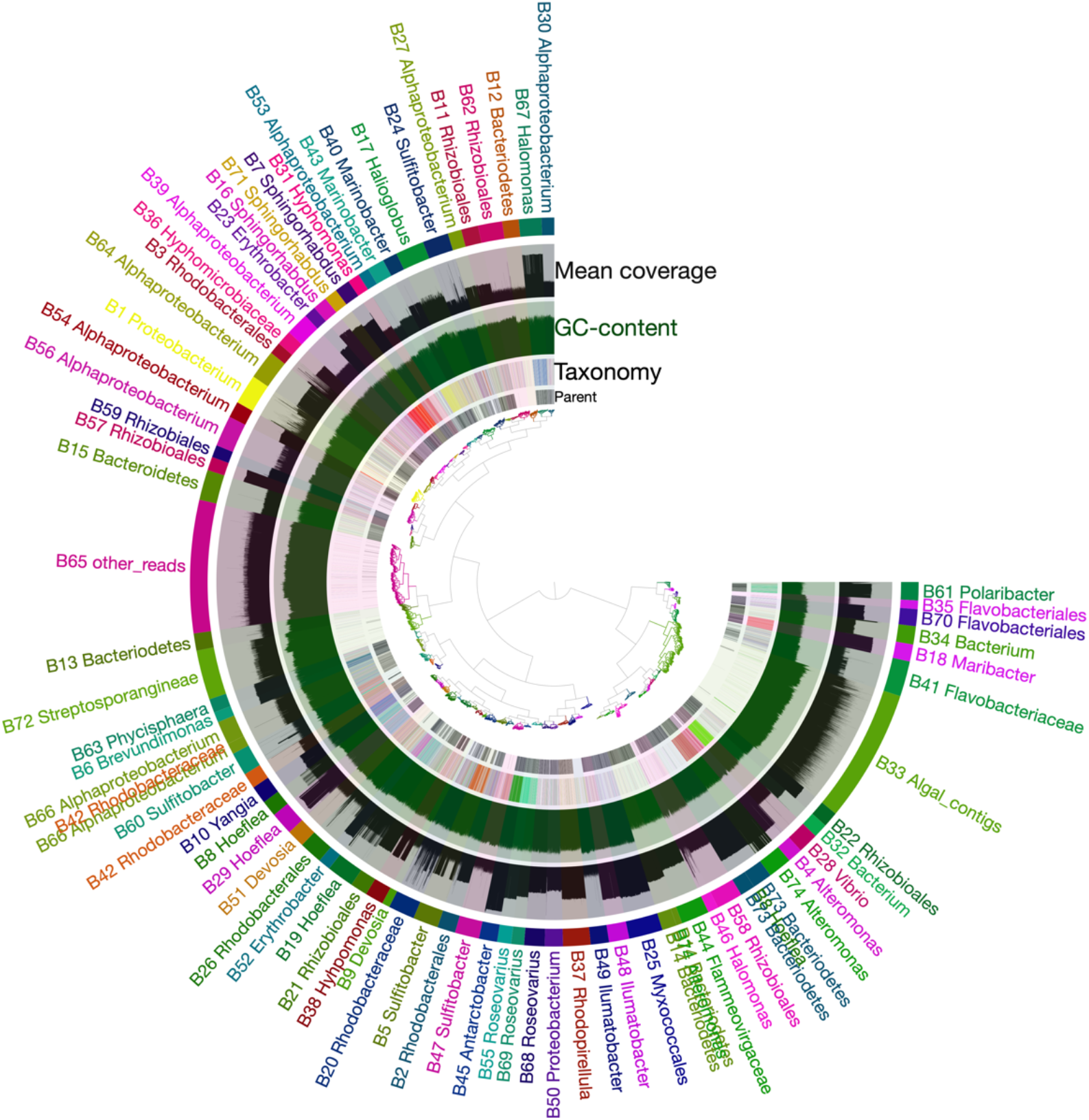
Anvi’o binning of assembled metagenome reads from a pool of all samples resulted in a total of 73 bacterial bins and one algal bin (B33). Contigs were clustered according to their kmer content, read coverage, and GC content, and manually grouped into bins. Colors were randomly assigned to the bins. For more information about each bin please refer to Supplementary Table S4.

### Algal gene expression

The PCA plots of algal gene expression show three groups (Figure 4A). The first group comprises the freshwater-tolerant holobionts H1 and H2 regardless of the salinity of their culture medium, and the other two groups correspond to the freshwater-intolerant holobiont in low (H3-15%) and full salinity (H3-100%), respectively. This global pattern was confirmed by DEG analyses carried out for each holobiont. H1 and H2 exhibited only a low number of differentially expressed genes (6 and 91, respectively; enriched in 0 and 6 GO categories, respectively), while in H3 2,355 genes (enriched in 100 GO terms) were differentially expressed (Figure 5A). A summary of the most important processes that were enriched among DEGs in each of the holobionts is given in Table 2, and details are provided in Supplementary Table S6. While H3 showed a more general response with several processes of primary metabolism being repressed in 15% NSW, H1 and H2 exhibited more specific responses. Interestingly processes classically associated with stress response, such as the production of heat shock proteins, were repressed in H3 low salinity.

**Table 2:** Summary of the differentially regulated processes during the response to low salinity in the algal host of each holobiont based on the summary of enriched GO terms from Supplementary Table S6. Upregulated genes are significantly induced in 15% NSW compared to 100% NSW and/or in holobiont H3 compared to holobiont 1+2 (interaction term). Downregulated refers to genes significantly repressed in the same conditions.

|  | Upregulated in 100% or H3 | Downregulated in 100% or H3 |
| --- | --- | --- |
| **Holobiont H1**<br>100% NSW vs.<br>15% NSW |  | heat shock protein<br>SAM-dependent methyltransferase<br>dynein heavy chain protein |
| **Holobiont H2**<br>100% NSW vs.<br>15% NSW | LHCR/LHCF<br>Cytochrome / PSII (cp)<br>Ribosomal proteins (cp)<br>Transcription / Translation | C5-epimerase<br>heat shock protein 70 |
| **Holobiont H3**<br>100% NSW vs.<br>15% NSW | Ammonium transmembrane transport<br>Mannose synthesis | Nitrate assimilation<br>Photosynthesis<br>Amino acid metabolism<br>Vitamin biosynthesis<br>Lipid metabolism |
| **Holobiont H1+H2**<br>vs.<br>**Holobiont H3** | Photosynthesis<br>Transmembrane transport<br>Transcription / translation<br>Amino acid metabolism<br>Lipid metabolism<br>Carbohydrate metabolism Nitrogen metabolism | Cytoskeleton |
| **Interaction term** | Transmembrane transport | Amino acid metabolism<br>Lipid metabolism<br>Photosynthesis<br>Vitamins<br>Carbohydrate metabolism |

**Figure 4:**
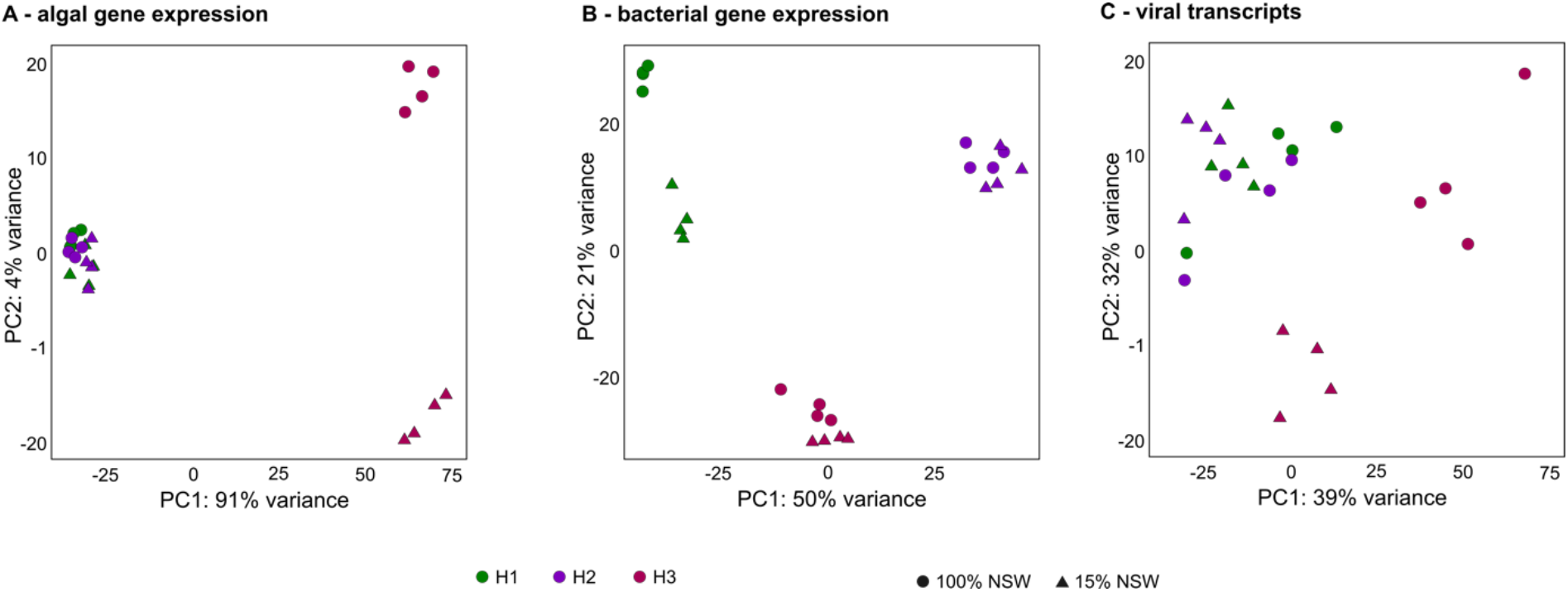
A) Principal component analysis (PCA) of algal gene expression. B) PCA of expression of bacterial reactions. C) PCA of viral transcripts

**Figure 5:**
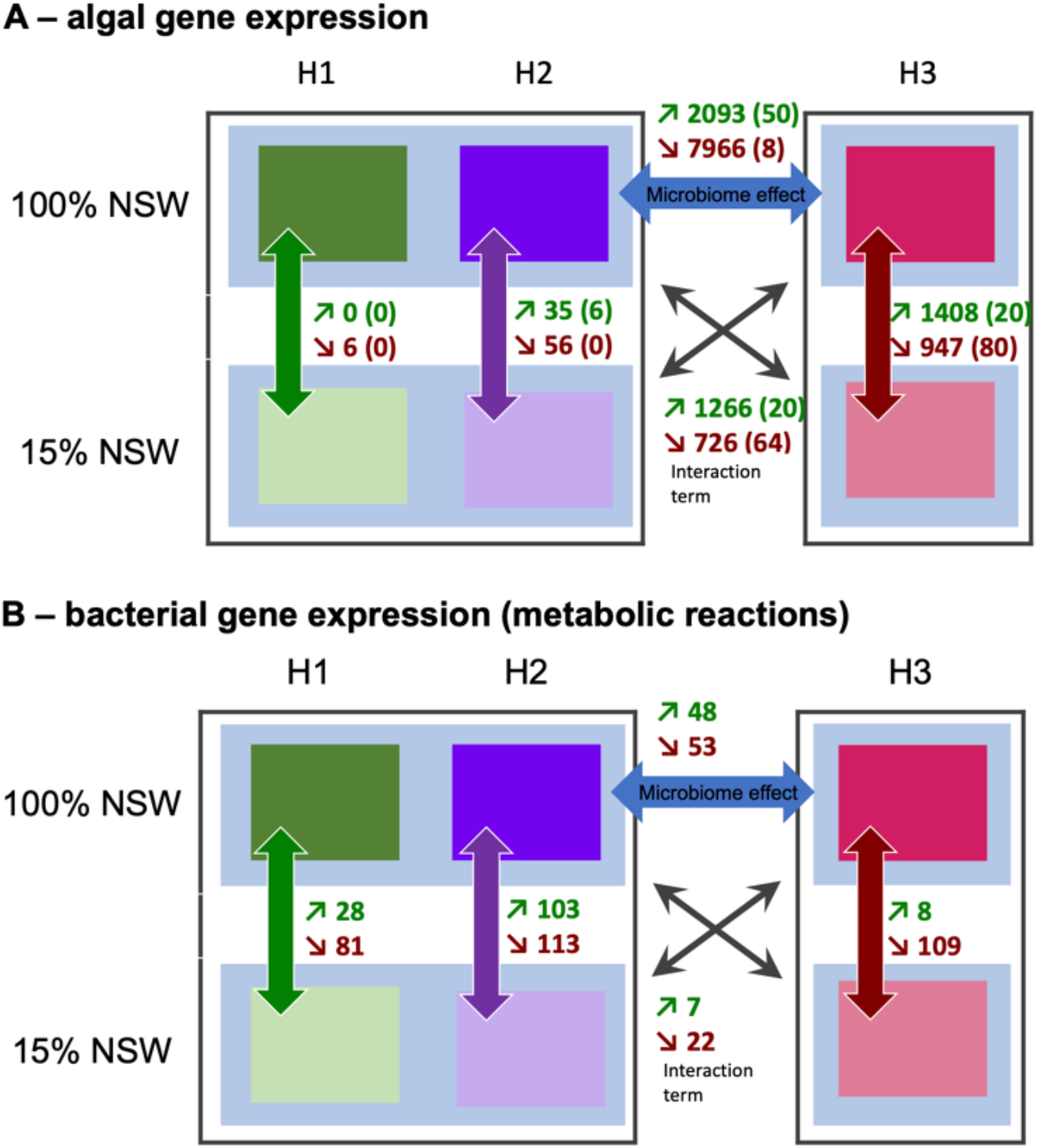
A) Differentially expressed algal genes. The figure shows the number of differentially expressed algal genes in each of the holobionts in 15% NSW compared to 100% NSW (H1, H2, and H3); H1 + H2 jointly compared to H3 in 100% NSW (microbiome effect), and the difference in the low salinity-response of H3 compared to that of H1+ H2 (interaction term, crossed arrows). Numbers in parentheses correspond to the number of overrepresented GO terms associated with the differentially regulated genes. B) Similar analysis as A, but on the bacterial transcriptome; in this case, the analysis was based on differentially expressed metabolic reactions rather than genes.

A key objective of this work was to determine how the freshwater-intolerant holobiont (H3) differed from the freshwater-tolerant holobionts (H1+H2), both in terms of basal gene expression in 100% NSW and regarding its response to low salinity. This was accomplished by DEG analysis of the data grouping H1+H2 and using a two-factor model (holobiont*salinity). In this model, genes significant for the factor holobiont correspond to basal differences in gene expression between freshwater-tolerant and freshwater-intolerant algae. Overall, 10,059 genes fell in this category; 7,966 were down-regulated, and 2,093 were upregulated in H3. Overrepresented GO terms associated with these genes cover a wide range of primary metabolic and cellular processes and are listed in Table 2 and, in more detail, in Supplementary Table S6.

In the same model, genes significant for the interaction term correspond to holobiont-specific gene regulation differences in response to low salinity. Here 1,266 genes were upregulated explicitly in H3 low salinity, and 726 specifically down-regulated. The physiological processes and GO terms overrepresented among these genes are summarized in Table 2 and listed in Supplementary Table S6. They comprise several typical responses to changing salinity, such as transmembrane transport (overrepresented in both up- and down-regulated genes), or lipid metabolism with activation of lipid breakdown in H3 in low salinity.

### Microbial gene expression

For the analysis of microbial gene expression patterns, despite the availability of a metagenome, a classical gene by gene and organism by organism analysis was not feasible due to the low final read coverage and the high number of microbes/microbial genes present (only 0.7% of all reads mapped to the 73 microbial bins). Therefore, we used two alternative approaches to exploit the available data. First, we summarized gene expression data for all genes within a given metagenomic bin to determine each bin’s overall transcriptomic activity. Secondly, we merged gene expression data for all genes predicted to catalyze the same metabolic reaction across all bins. These latter data were used to examine differences in the overall metabolic activity of the entire microbiome in the tested conditions.

Normalized transcriptional activity per bin was visualized in a heat map (Figure 6). It shows that each of the three holobionts is characterized by the transcriptomic activity of different metagenomic bins, and smaller differences within each of these clusters separate the 100% NSW from the 15% NSW conditions. A similar pattern was also observed in the PCA plot based on the microbiome’s overall metabolic activity: again, holobiont was the main separating factor, but within each holobiont, separation according to the salinity treatment is visible (Figure 4B).

**Figure 6:**
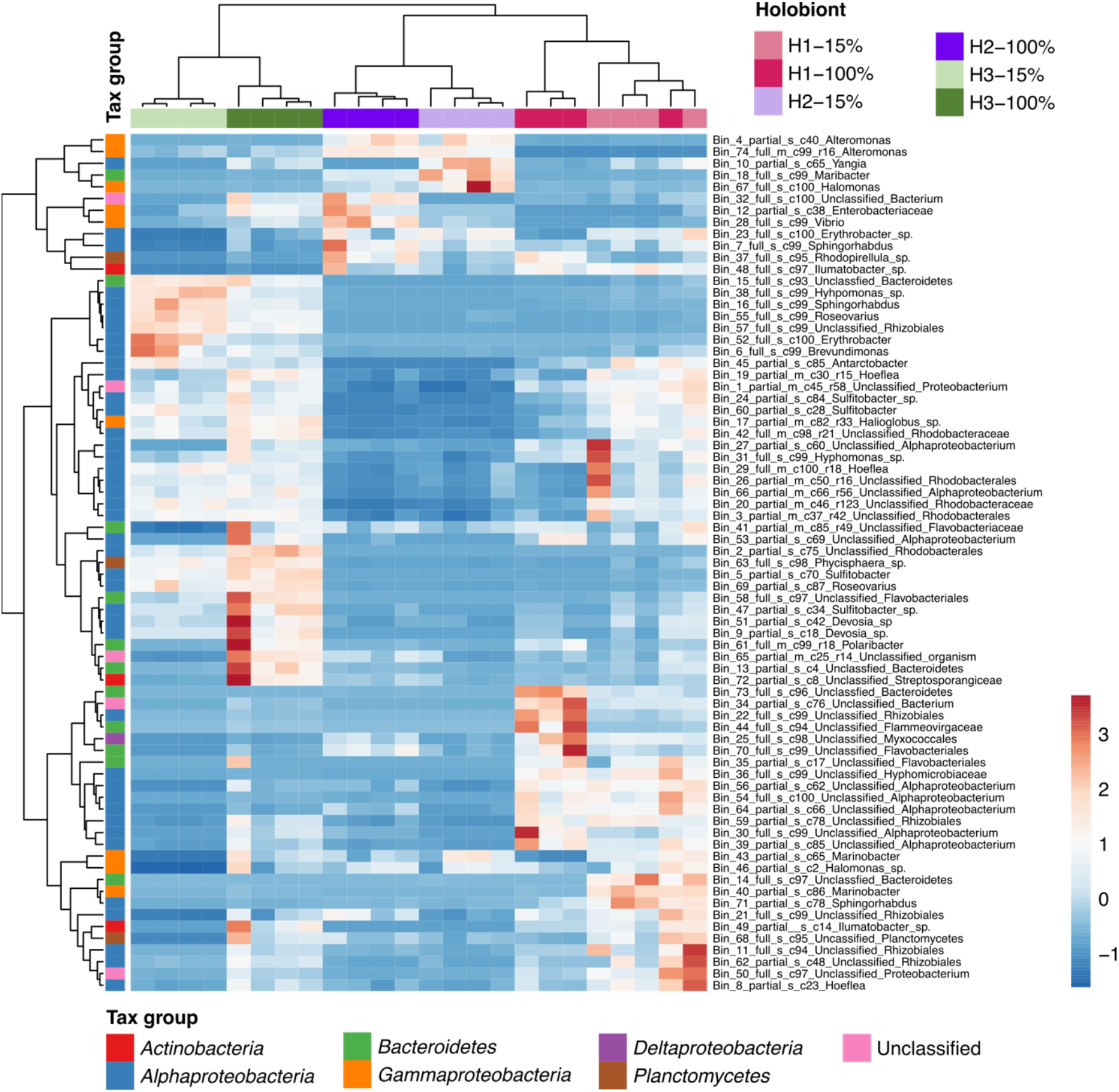
Heatmap based on hierarchical clustering of normalized transcriptional activity per bacterial bin in each holobiont and condition (Pearson correlation coefficient; clustering method: average linkage; unit variance scaling of row data). Red indicates high relative transcriptomic activity in the given condition, blue low activity. “15%” corresponds to the treatment with 15% natural seawater (NSW), “100%” to the treatment with 100% NSW. Bins labeled “full” are predicted ≥ 90% complete, bins labeled “partial” <90%. The predicted completeness in % is given after the “c” in the bin name.

To determine the metabolic specificities of the bacteria associated with the tested holobionts and conditions, expression data for each of the 3,957 metabolic reactions (Supplementary Table S5) were subjected to differential expression analysis. Hundred nine significant reactions were identified in holobiont H1, 226 in holobiont H2, and 117 in holobiont H3 (Figure 5B, Supplementary Table S7). In the microbiome of holobiont H1, gene expression differences between the salinity levels concerned most importantly glycine biosynthesis (induced in low salinity) and quinone biosynthesis as well as quorum sensing (repressed in low salinity). In holobiont H2, glycine, sorbitol, butanol, and polyamine metabolism were induced, and several genes related to nucleotide degradation and quinone metabolism were repressed in low salinity (among other reactions). In holobiont H3, unlike in holobiont H1 and H2, osmolyte production (glycine-betaine, ectoine) was repressed in low salinity, along with some carbohydrate degradation pathways and ATP as well as NAD metabolism. No pathways in this holobiont contained >50% of induced reactions in low salinity (see Supplementary Table S7) for a complete list of reactions). Only one reaction, catalyzed by a diaminobutyrate aminotransferase (R101-RXN), was repressed in low salinity by all holobionts and no reactions were significantly upregulated in all holobionts. Comparing global expression patterns of holobiont H3 with holobionts H1 and H2 in full salinity highlighted only 4 pathways: hydrogen oxidation, phosphonoacetate degradation, pyruvate fermentation, and hydrogen to fumarate electron transfer, all of which were upregulated in holobiont H3.

Examining reactions significant for the interaction term, i.e. microbial reactions for which the response to low salinity differed between low-salinity tolerant and intolerant holobionts, only one pathway, autoinducer AI-1 biosynthesis, emerged. This pathway was explicitly upregulated in the fresh water-intolerant holobiont H3 in low salinity conditions, notably by bacteria of the genera *Hoeflea* (bin 29), *Roseovarius* (bins 55 and 69), and *Sulfitobacter* (bin 5). The other 21 significant reactions did not constitute pathways with >50% of genes regulated, but, given their potential importance, they were manually grouped into 8 metabolic categories: ectoine synthesis, phospholipids, phosphate metabolism, selenate reduction, seleno-amino acid biosynthesis, carbon metabolism, vitamins, and DNA repair, all of which were repressed or absent in holobiont H3 in low salinity (Table 3). Notably, the demethylphylloquinone reduction reaction (RXN-17007) constituting the last step of the biosynthetic pathway of vitamin K was expressed by three bins, although at very low levels. Bin 32 (unclassified bacterium) expressed this reaction in holobionts 2 and 3 in seawater, bin 19 (*Hoeflea*) expressed the reaction in holobiont 1 (both seawater and 15% NSW conditions) and holobiont 3 in seawater, and bin 67 (*Halomonas*) expressed the reaction in holobionts 2 in 15% NSW. However, none of the bacteria in our metagenome expressed RXN-17007 in H3 in 15% NSW.

**Table 3:** Metabolic reactions with different low salinity responses in H3 compared to H1 and H2 (interaction term). The first column contains the MetaCyc reaction ID and, in parentheses, the corresponding enzyme. If the same enzymes carried out several (similar) reactions, these were grouped and only one representative is shown. If different than 1, the total number of reactions catalyzed by this enzyme is given in parentheses following an x after the enzyme name. “Induced” means that this reaction was more strongly upregulated in H3 in response to 15% NSW than in H1 and H2. Repressed or absent means the reaction was downregulated or absent in H3 in 15% NSW.

|  | Regulation | Important taxa |
| --- | --- | --- |
| **Quorum sensing** |  |  |
| 2.3.1.184-RXN (acyl-homoserine-lactone synthase) (x6) | Induced | *Sulfitobacter*, *Roseovarius*, *Hoeflea* |
| **Ectoine synthesis** |  |  |
| R101-RXN (diaminobutyrate aminotransferase) | Repressed | *Antarctobacter*, *Roseovarius*, *Halomonas* |
| R102-RXN (diaminobutanoate acetyltransferase) | Repressed | *Antarctobacter*, *Roseovarius*, *Halomonas* |
| R103-RXN (ectoine synthase) | Repressed | *Antarctobacter*, *Roseovarius*, *Halomonas* |
| **Phospholipids** |  |  |
| GLYCPDIESTER-RXN (glycerophosphoryl diester phosphodiesterase) (x3) | Repressed | *Antarctobacter*, *Alteromonas*, *Sphingorhabdus*, *Sulfitobacter*, *Erythrobacter*, uncl. *Rhizobiales* |
| LIPIDXSYNTHESIS-RXN (UDP-2,3-diacylglucosamine diphosphatase) | Repressed | *Hoeflea*, *Phycisphaera* sp., uncl. *Bacteroidetes* |
| **Phosphate turnover/metabolism** |  |  |
| 3.11.1.2-RXN (phosphonoacetate hydrolase) | Repressed | *Hoeflea* |
| ABC-27-RXN (phosphate ABC transporter) | Repressed | *Sphingorhabdus*, *Rhodobacteraceae*, *Hoeflea*, *Antarctobacter*, *Erythrobacter*, *Rhizobiales*, *Phycisphaera*, *Alteromonas* |
| RXN0-1401 (ribose 1,5-bisphosphate phosphokinase) | Repressed | *Halomonas*, *Hoeflea*, uncl. *Rhizobiales*, *Antarctobacter* |
| **Selenate reduction/seleno-amino acid biosynthesis** |  |  |
| RXN-12720 (ATP-sulfurylase) (x2) | Repressed | *Illumatobacter*, uncl. bacterium, uncl. *Bacteroidetes*, *Alteromonas* |
| RXN-12730 (5-methyltetrahydropteroyltri-glutamate--homocysteine S-methyltransferase) | Repressed | *Halomonas* |
| **Carbon metabolism** |  |  |
| RXN-961 (ribulose biphosphate carboxylase/oxygenase) | Repressed | Uncl. bacterium, *Algoriphagus*, *Lutibacter* |
| 2.8.3.22-RXN (Succinyl-CoA--L-malate CoA-transferase) (x4) | Repressed | *Sphingorhabdus*, *Hoeflea*, uncl. *Rhodobacterales*, *Hyphomonas* |
| RXN-14365 (D-psicose 3-epimerase.) | Repressed | *Hoeflea*, *Antarctobacter* |
| **Vitamins** |  |  |
| RXN-17007 (Demethylphyloquinone reductase) | Absent | *Hoeflea*, *Halomonas*, uncl. bacterium |
| **DNA repair** |  |  |
| 3.1.21.2-RXN (Endonuclease) | Repressed | *Hoeflea* |

### Expression of viral sequences

The analysis of viral sequences in the metatranscriptome revealed marked differences in viral abundances depending both on the holobiont and the culture condition (Figure 4C). The global patterns followed that of the algal transcriptome data, with holobiont 3 grouping apart from holobionts 1 and 2, and with a clear difference in viral composition in holobiont 3, depending on salinity. Holobiont 3 in 100% salinity conditions exhibited the highest relative abundance of viral sequences (Figure 7). The majority of annotated viral reads belonged to the *Phycodnaviridae*, dsDNA viruses known to infect algae (Wilson *et al.*, 2009). They were present in all samples and most abundant in H3-100%. The next most abundant groups of viruses were DNA phages belonging to the *Microviridae* and *Caudovirales*. These groups specialize in bacterial hosts (King *et al.*, 2012; Doore and Fane, 2016). *Retroviridae* occurred specifically in holobiont 2 in 100% salinity, and of *Alloherpesviridae* also in holobiont 2 in 100% salinity as well as in one replicate of H3-15%.

**Figure 7:**
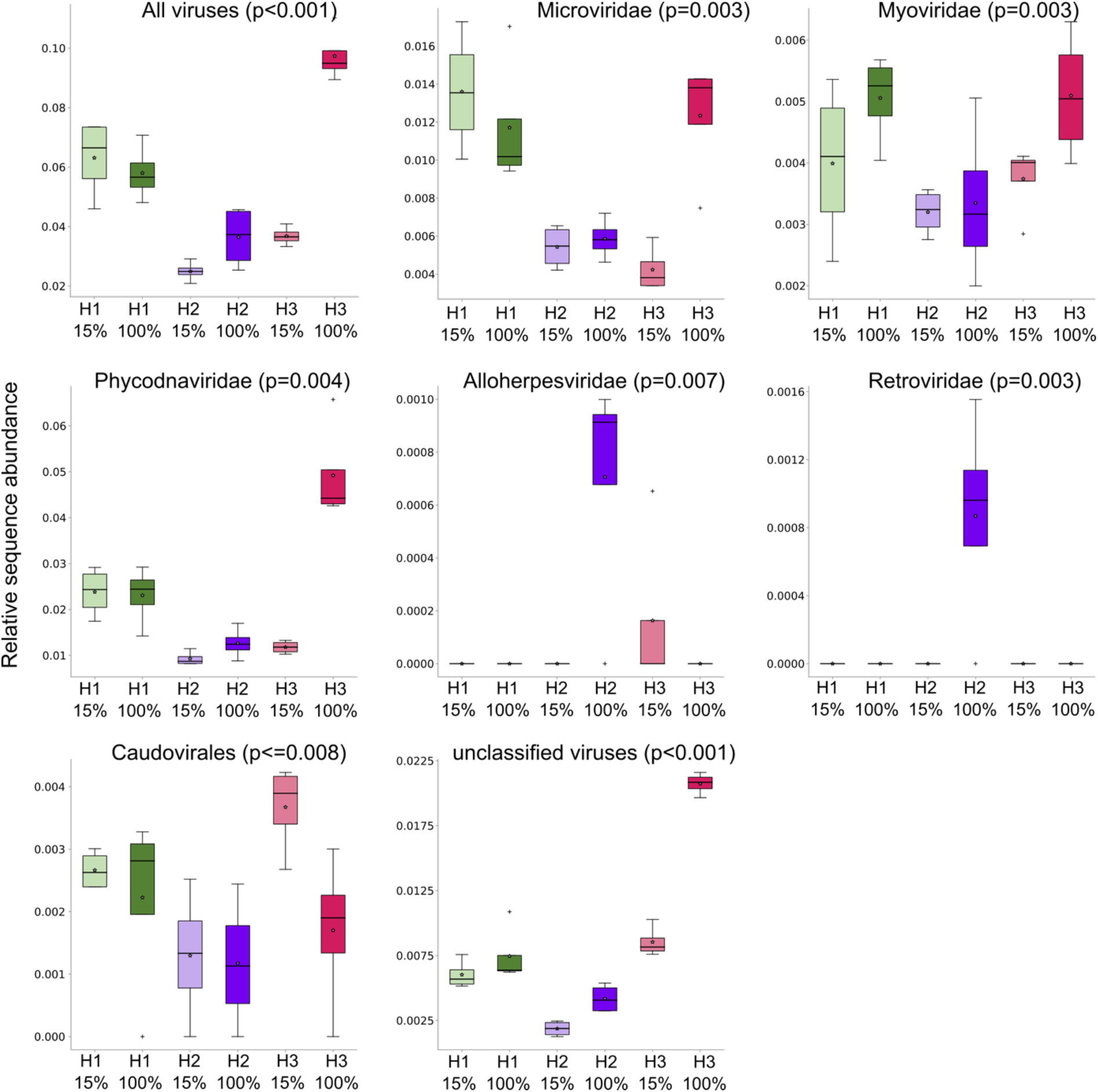
Box plot of relative abundances of viral sequences belonging to different families in all samples (H1-H3: holobiont 1-3). Values are given in % of the total number of RNAseq reads for each sample. P-values correspond to the results of an ANOVA across all six conditions (4 replicates each). 15% corresponds to the treatment with 15% NSW, 100% to the treatment with 100% NSW.

### Metabolite profiling of algal holobionts

The metabolite dataset contained 609 features that were (after normalization) used for statistical testing. In total, 72 features were significantly different in at least one of the tested conditions (Supplementary Table S8). Differences between H1 and H2 were negligible, with only one feature differing between the low and high salinity conditions of these two holobionts (a peak putatively corresponding to histidinol in H1, and a peak putatively corresponding to hydroquinone in H2 both down-regulated in low salinity). This was confirmed by cluster analysis (Figure 8), which does not separate these conditions. H3, however, exhibited differences in metabolic profiles depending on the salinity with 4 upregulated (quinic acid, 5-propionate-hydantoin, and two unknown features) and 35 down-regulated features in low salinity, including a feature annotated as phytol (peak #145, a precursor of vitamin K) and several primary metabolites such as galactoglycerol, glycine, alanine, valine, oxoproline. For all of these, the regulation was specific to H3. Furthermore, even under control conditions (100% NSW), metabolite profiles of H3 differed from holobionts H1 and H2, exhibiting higher abundances of 18 features in H3 compared to H1+H2 (including glycerol 3-P, pentafuranose, putrescin), and a lower abundance of 29 features (including alanine, galactoglycerol, and valine; see Figure 8).

**Figure 8:**
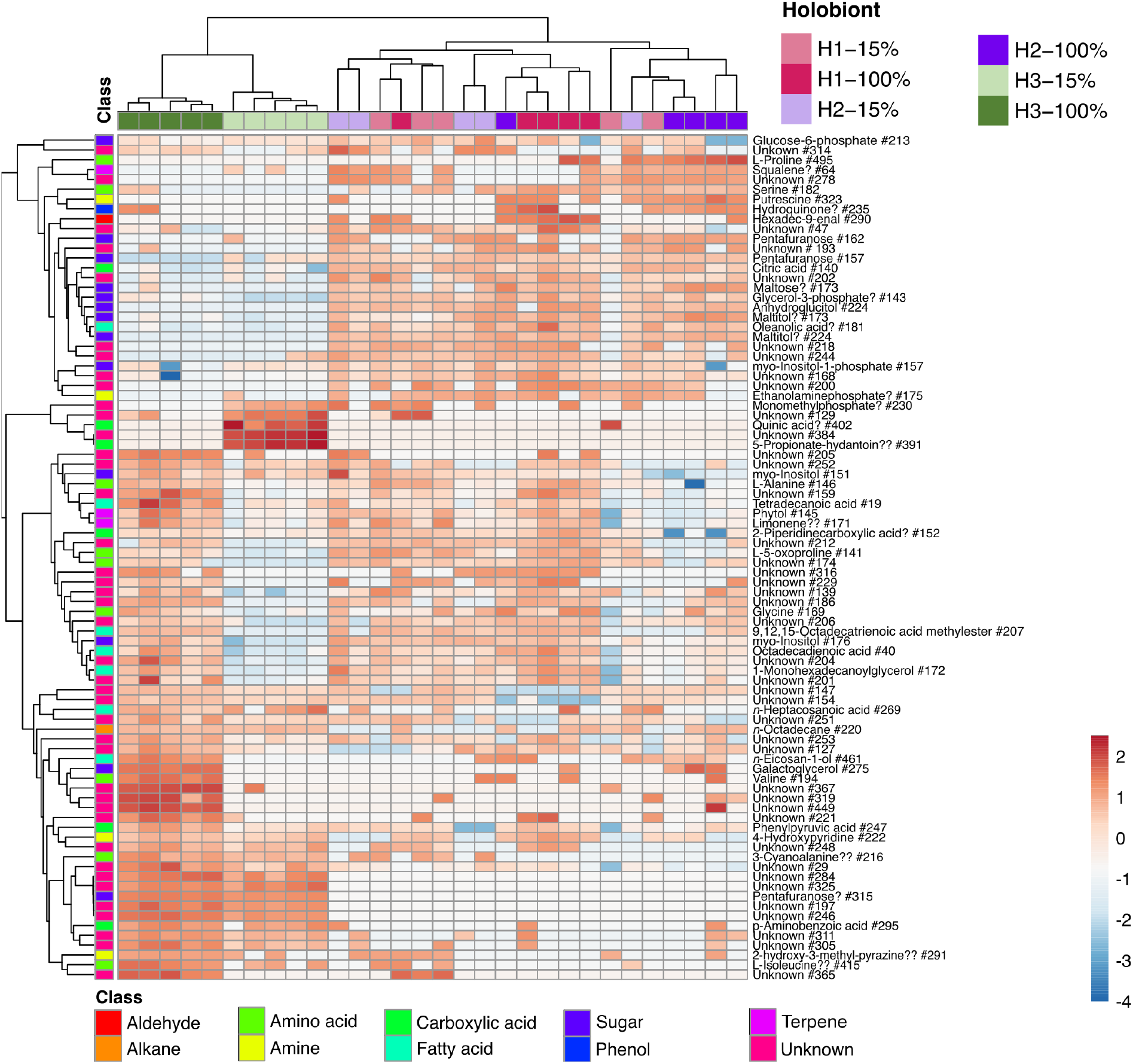
Heat map based on the abundance of each of the metabolites tested as significant in at least one condition (Pearson correlation coefficient; clustering method: average linkage; unit variance scaling of row data). 15% corresponds to the treatment with 15% NSW, 100% to the treatment with 100% NSW. “?” indicates features with reverse match score (R)<800, “??” features with R<700. Features were labeled “unknown” for R<600.

## Discussion

This study aimed to identify bacteria and bacterial functions that could support *E. subulatus* holobionts to acclimate to low salinity. Three algal-bacterial holobionts that differed in their capacity to grow in low salinities were created. Our data show that variations in microbiome impacted algal gene expression profiles and metabolomic features. Low salinity-tolerant holobionts were those that were given fewer antibiotics and had fewer differentially expressed genes and metabolites in response to changes in salinity. Our findings show that low salinity treatments were not particularly stressful for the algal host in freshwater-tolerant holobionts, despite microbiome changes resulting from the treatment. In the fresh water-intolerant holobiont, on the other hand, basal expression levels, metabolite profiles, and even viral expression levels were altered, and we observed a robust host response to salinity changes, similar to that described by Dittami *et al.* (2012). Algal holobionts were allowed to recover in antibiotic-free natural seawater medium (NSW) for weeks after the initial antibiotic treatment in our experiments. We, therefore, assume that the differences in the algal response to low salinity are related mainly to differences in the microbiome. This assumption is in line with the results of inoculation experiments demonstrating that low salinity tolerance in antibiotic-treated cultures has previously been restored by restoring the microbiome (Dittami *et al.*, 2016). Our data also suggest that viral transcription changes may further complexify this already complex system, as the relative abundance of viral reads varied depending on the holobiont and culture condition. These variations affected both viruses likely to infect the host and the associated bacteria. Such tri-partite trans-kingdom interactions have been described in mammalian guts (Pfeiffer and Virgin, 2016), but have, to our knowledge, not been considered in macroalgal research so far.

### Three scenarios of microbial impacts on host stress tolerance

Here we discuss the generated metatranscriptomic, metagenomic, and metabolomic data in the light of three different scenarios of how bacteria may impact algal low salinity tolerance:

1. Members of the microbiome of low salinity-tolerant holobionts may directly produce compounds such as osmolytes or chaperones that enhance algal low salinity tolerance or stimulate the alga to do so.
2. The microbiome provides essential services to the alga regardless of the salinity, but under stress, the microbiome of the low salinity-sensitive holobiont may no longer provide these services, ultimately leading to a reduction/stop in growth.
3. Under stressful conditions, the equilibrium in the microbiota may be disrupted, and certain microbes may proliferate and become harmful to the host. This phenomenon, termed dysbiosis, can be triggered both by algal and bacterial signals and possibly mitigated by the presence of other microbes or viruses.

### No clear signs of microbial contributions to algal stress response

In the first scenario, the alga-associated microbiome is assumed to produce compounds that enhance algal stress tolerance actively. In terrestrial environments, studies have highlighted, for instance, the microbial production of plant hormones, which activate plant defenses, making them more resistant against pathogens, or the uptake of nutrients (e.g. via the production of siderophores, that enhance the host ability to survive in low-nutrient environments) (Numan *et al.*, 2018). When examining the low salinity response of the different holobionts, we found little difference in algal gene expression or the examined metabolite profiles in the low-salinity tolerant holobionts. Furthermore, none of the observed changes corresponded to the induction of classical stress response genes.

On the bacterial side, the only low-salinity induced microbial pathway common to both low salinity-tolerant holobionts was glycine synthesis, but we did not detect significant changes in glycine concentrations in the tissues of these holobionts. Glycine has been shown to enhance growth in some microalgae, including diatoms (Berland *et al.*, 1979). However, no data is available on the impact of external glycine on brown algal growth rates. Furthermore, *Ectocarpus* can synthesize glycine and does so primarily during the daytime (Gravot *et al.*, 2010). This does not exclude a role of glycine or other bacterial compounds in the brown algal stress response, but our transcriptomic data provide little support for this hypothesis.

An alternative way microbes may increase host tolerance to stressors is by priming the host and activating its “defenses” even before exposure to stress – a process previously reported in kelps (Thomas *et al.*, 2011). Indeed, a comparison of transcriptomic and metabolic profiles of holobionts H1 and H2 vs H3 revealed fundamental differences, indicating that the host compartments of the different holobionts were not in the same physiological state at the start of the experiments. However, most of the processes upregulated in H1+H2 were related to the cytoskeleton and not GO categories such as “response to stimulus” or more specific sub-categories. Thus, while our results cannot exclude potential defense priming effects, the algal host’s transcriptomic regulation does not support this hypothesis.

### Loss of microbial services in non-tolerant holobionts: vitamin K

A second scenario is that changes in the microbiome triggered by the salinity change result in a lack of microbial services essential for the alga. Bacteria are known to provide, e.g. growth hormones to brown algae, including *Ectocarpus* (Pedersén, 1973; Tapia *et al.*, 2016), and more profound metabolic interdependencies have been predicted based on metabolic networks (Burgunter-Delamare *et al.*, 2020). However, an exhaustive list of these microbial contributions is still missing. A loss of such benefits could result from shifts in the microbiome composition, changes in the activity of different microbes, or specific changes in bacterial gene expression patterns. Our transcriptomic and metabolomic data show that holobiont H3 had fundamentally different basal profiles even in seawater before applying any low salinity stress. Notably, several primary metabolic processes were activated compared to holobionts 1 and 2, such as the synthesis of amino acids and lipids, photosynthesis, carbohydrate metabolism, and transcription/translation. In contrast, several primary metabolites were less abundant (proline, alanine, glycine, serine, citric acid, and more, Figure 7). One possible interpretation of these observations is that the host in holobiont H3 was able to compensate for the absence of bacterial functions as long as it was growing in seawater but was no longer able to do so when transferred to low salinity, resulting in a repression of primary metabolism and a reduction of growth.

Our transcriptomic data highlighted bacterial metabolic processes that were less expressed and possibly absent specifically in H3 in low salinity (Table 3). Based on the *Ectocarpus* genome (Cock *et al.*, 2010), most of these processes are likely to be achievable by the algal host itself without requiring input from bacteria (carbon, phosphate, selenate, and phospholipid metabolism, DNA repair). While this does not preclude that these compounds or activities may be helpful for the host, their provision from external sources is likely not essential.

In the same vein, the synthesis of ectoine was repressed in the bacterial metatranscriptome H3 in low salinity. Ectoine is known to serve as an osmolyte in bacteria (Czech *et al.*, 2018) and further studies are necessary to determine if ectoine might be also released to the environment to the benefit of the host. Microalgae without an associated microbiome contain ectoine in small amounts, pointing towards a dual origin of this metabolite in the algae from their own biosynthesis as well as from uptake (Fenizia *et al.*, 2020). However, ectoine might be only used as an osmolyte for bacteria in 100% NSW, and may no longer be required at low salinity.

Vitamin K is involved in the functioning of the photosystem in land plants. Mutants of *Arabidopsis* and *Cyanobacteria* missing the last reaction of its biosynthetic pathway are viable but exhibit increased photosensitivity (Fatihi *et al.*, 2015). In *E. subulatus*, just as in *E. siliculosus, Saccharina latissima*, and *Cladosiphon okamuranus* (Nègre *et al.*, 2019; Dittami, Corre, *et al.*, 2020), the last step of the vitamin K biosynthetic pathway is absent from the algal metabolic network. It suggests that these algae cannot produce vitamin K independently, although this compound has previously been detected in kelps (Yu *et al.*, 2018). If, as suggested by our transcriptomic and metabolic data, bacterial vitamin K production is repressed specifically in H3 in low salinity, it is plausible that the resulting lack of vitamin K may negatively impact algal growth. This constitutes a promising hypothesis to be tested, for instance via complementation experiments.

### Indications for dysbiosis

The last of the three scenarios assumes that, in the fresh water intolerant holobiont, a change in salinity led to a change in the bacterial community or activity, notably the propagation of specific microbial strains that harm the host (dysbiosis). Dysbiosis is a well-studied phenomenon, especially in mammalian models. It is known to be the basis of several diseases (e.g. Hawrelak and Myers 2004) and assumed to be widespread also in marine environments (Egan and Gardiner, 2016).

In our data, we observed a significant induction of the AI-1 pathway, specifically in H3 in low salinity (i.e. in the interaction term), supporting the hypothesis of the activation of quorum sensing compounds. Quorum sensing (QS) and AI-1, in particular, have been linked to dysbiosis in several model systems. For instance, in the coral *Pocillopora*, AI-1 type QS compounds have been related to coral bleaching (Zhou *et al.*, 2020). Similarly, in *Acropora*, inhibitors of AI-1 have been shown to prevent white band disease (Certner and Vollmer, 2018). In several microalgae, QS molecules have also been associated with the bacterial production of algicidal proteins or compounds such as proteases, amylases, quinones, or in many cases, unknown compounds (Paul and Pohnert, 2011; Demuez *et al.*, 2015). Furthermore, we observed a 4-fold increase in the relative bacterial ‘activity/abundance’ as measured by the ratio of bacterial mRNA to algal mRNA reads, suggesting strong bacterial proliferation as typical for dysbiosis. This increase in transcription levels concerned primarily bacteria that were little active in the other conditions (unclassified *Bacteroidetes*, *Hyphomonas*, *Sphingorhabdus*, *Roseovarius*, unclassified *Rhizobiales*, *Erythrobacter*, *Brevundimonas*, Figure 6).

Both observations, the induction of QS genes and the increase in bacterial abundance, fit well with the scenario of dysbiosis. However, there are two significant limitations: First, QS compounds are not exclusively linked to dysbiosis and virulence – they may also be involved in processes such as the regulation of bacterial motility, biofilm formation, or the regulation of nutrient uptake (Zhou *et al.*, 2016), and in our data, we did not observe the induction of virulence-related genes. This could be explained by the nature of our analyses, which considers only genes with known functions that are represented in the MetaCyc database. Any (unknown) compounds or genes not represented in MetaCyc would not have been detected. The second limitation is linked to the first. Without further targeted experiments, it is not possible to assert if the induction of QS genes may be an indirect cause of the poor algal physiological state, or rather if the behavioral response of the bacteria to this compound causes poor algal health. Based on these observations, we believe that the role of QS during freshwater acclimation may be of interest for future studies, e.g. with QS inhibitors, and these studies should also specifically examine the production of potential virulence factors.

## Conclusion

The case of the freshwater strain of *E. subulatus* and its reliance on its microbiome for growth and survival in freshwater is an excellent example of the importance of host-symbiont interactions. The system also demonstrates the difficulties that arise when attempting to understand the exact interactions in such complex systems, especially when they are not well-established models: fully controlled experiments to elucidate the specific functions are not (yet) possible because we have not (yet) been able to cultivate or identify the right mix of bacteria required for freshwater tolerance. As a result, we chose to empirically modify holobiont composition and then study its behavior using a combination of metabolomics, metatranscriptomics, and metagenomics. Metabolic networks were used as a filter to combine and interpret the resulting complex datasets. This data only covers a portion of the biology of the organisms studied, but in our case suggested potential metabolic roles of associated bacteria in the supply of vitamin K, as well as a possible role of quorum sensing compounds in the holobionts that no longer exhibited growth - both hypotheses that can now be tested in targeted experiments, such as by supplementing vitamin K to the culture medium or by using inhibitors of quorum sensing. Lastly, our findings highlight the importance of considering viruses as a potential factor influencing holobiont acclimation to environmental change by interacting with both the host and the bacterial microbiome.

## Acknowledgments

We thank Laurence Dartevelle for advice on algal culturing; Sylvie Rousvoal for advice on DNA and RNA extractions; David Green for advice on the metagenome analysis; Claire Gachon for project coordination and helpful discussions; and the Institut Français de Bioinformatique (ANR-11-INBS-0013), the Roscoff Bioinformatics platform ABiMS (http://abims.sb-roscoff.fr/), as well as the Genouest platform (https://www.genouest.org/) for computing and storage resources. This work has received funding from the European Union’s Horizon 2020 research and innovation program under the Marie Sklodowska-Curie grant agreement number 624575 (ALFF, to HK, GC, TW, SD), the ANR project IDEALG (ANR-10-BTBR-04) “Investissements d’Avenir, Biotechnologies-Bioressources”, the CNRS momentum call, the German Research Foundation through the CRC 1127/2 ChemBioSys – 239748522 grant (to GC, TW), and Conseil Departemental 29 via the VIRALG project (to EK, SD).

